# Stimulation of cholesterol efflux inhibits herpesvirus nuclear egress

**DOI:** 10.1101/2025.10.03.680238

**Authors:** Eric S. Pringle, Carolyn-Ann Robinson, Alexa N. Wilson, Ethan C. M. Thomas, Janani Krishnan, Andrea L-A. Monjo, Katrina Bouzanis, Bruce W. Banfield, Brett A. Duguay, Craig McCormick

## Abstract

Cholesterol in herpesvirus envelopes and host membranes supports membrane fusion during virus entry. However, little is known about how cholesterol affects other aspects of the viral replication cycle. Here, using an infection model that begins with reactivation from latency, bypassing the entry step, we demonstrate that depleting intracellular cholesterol inhibits the production of infectious Kaposi’s sarcoma-associated herpesvirus (KSHV). Treatment of latently infected cells with the liver X receptor α (LXRα) agonist 22(*R*)-hydroxycholesterol (22OH) increased expression of sterol response genes, including the cholesterol efflux pump ATP binding cassette subfamily A member 1 (ABCA1), and reduced intracellular cholesterol. 22OH treatment initially enhanced lytic reactivation, but diminished viral protein accumulation during late replication, while altering accumulation of host proteins that regulate the cell cycle and innate immunity. Cholesterol efflux led to an increase in the proportion of capsids lacking viral genomes in the nucleus and a reduction in nucleocapsids accessing the cytoplasm. Next, we used a herpes simplex virus (HSV) infection model to explore if 22OH affected the conserved herpesvirus nuclear egress complex (NEC), comprised of two viral proteins that accumulate at the inner nuclear membrane and facilitate primary envelopment. We found that 22OH inhibited HSV-1 and HSV-2 replication and spread in plaque reduction assays and impaired localization of NEC proteins pUL31 and pUL34 to the nuclear periphery. These findings indicate that cholesterol efflux inhibits herpesvirus replication by interfering with the formation of nucleocapsids competent to complete nuclear egress.

**IMPORTANCE:** Cholesterol plays key roles in regulating the fluidity of cellular membranes and the trafficking and lateral diffusion of transmembrane proteins. Herpesviruses encode a variety of transmembrane proteins that traffic to different cellular compartments, while herpesvirus capsids and tegument proteins interact with nuclear and trans-Golgi network membranes throughout the primary and secondary envelopment process, respectively. This produces viral particles with cholesterol as an envelope component which is required to support membrane fusion during entry. Our discovery that cholesterol efflux inhibits KSHV nuclear egress suggests that cholesterol plays important supportive roles at this stage of lytic replication. Moreover, we observed that cholesterol efflux mediated by oxysterol treatment has widespread effects on host cell protein expression, resulting in a non-classical antiviral state. Cholesterol likely supports the proper trafficking of viral proteins to nuclear sites of virion assembly or is influencing the fluidity of nuclear membranes used for viral primary envelopment and de-envelopment during nuclear egress.

## INTRODUCTION

Kaposi’s sarcoma-associated herpesvirus (KSHV) is the infectious cause of Kaposi’s sarcoma (KS), primary effusion lymphoma (PEL) and multicentric Castleman’s disease (1–3). Like all herpesviruses, KSHV has a host-derived lipid envelope, studded with glycoproteins that govern virus tropism, entry and cell-to-cell spread. Virion envelopment occurs twice during the assembly process. First, following DNA packaging in the nucleus, capsids acquire a temporary lipid envelope from the inner nuclear membrane (INM) concomitant with nuclear egress. This process is facilitated by a nuclear egress complex (NEC), composed of two viral proteins that display high conservation amongst herpesviruses (4–12). Intensive study of alphaherpesvirus NECs has yielded insights into the structure and function of herpes simplex virus type 1 (HSV-1), herpes simplex virus type 2 (HSV-2) and pseudorabiesvirus (PRV) NEC complexes. During herpesvirus infection, the NEC recruits cellular and viral kinases, resulting in phosphorylation and solubilization of the nuclear lamina (7, 13–16). Similarly, the KSHV NEC, comprised of ORF67 and ORF69, causes nuclear envelope remodelling and disassembly of the nuclear lamina, allowing capsid budding from the INM (17). The mechanisms behind NEC recruitment of cellular and viral proteins remains incompletely understood. Following primary envelopment, enveloped particles fuse with the outer nuclear membrane (ONM), resulting in de-envelopment. De-enveloped capsids transit through the cytoplasm and acquire a final lipid envelope by budding into a post-Golgi compartment, likely the trans-Golgi network (TGN) or a late endosomal compartment, to access the host cell secretory pathway and complete egress (18).

Viruses modulate host lipid dynamics including lipid synthesis, metabolism and storage (19). For example, human cytomegalovirus (HCMV) infection results in increased glucose uptake and expression of genes involved in fatty acid synthesis (20–22). In HCMV infected cells, increased expression of a fatty acid elongase increases very long chain fatty acids (VLCFAs), which are incorporated into viral envelopes and required for assembly and virion infectivity (23, 24). Furthermore, lipids are altered during both KSHV latency and lytic replication. During latent KSHV infection of endothelial cells, fatty acid synthesis, cholesterol esterification and intracellular lipid droplets are increased and the inhibition of fatty acid synthesis during latency triggers cell death (25). Altering host cell lipid metabolism during KSHV latency may prime cells for efficient viral lytic replication. Reactivation from latency alters cellular lipid metabolism by increasing triglyceride synthesis and reorganizing intracellular membranes, where fatty acid synthesis is required to support KSHV virion maturation and egress (26).

Cholesterol is an integral component of cellular membranes that controls membrane fluidity, permeability, and organization and there is emerging evidence that cholesterol levels affect enveloped virus egress and the structural integrity of virions. Cell membrane cholesterol is required to assist membrane fusion events, initiated by cellular processes or viruses, either at the plasma membrane or when fusion occurs at endosomal membranes post-internalization (27–34). Cholesterol incorporation into viral envelopes during secondary envelopment at the trans-Golgi network (TGN) has been shown to affect virion stability, infectivity, and membrane fusion during viral entry (35–42). A study by Wudiri and Nicola showed that cholesterol depletion can negatively impact HSV-1 replication (40). Since cholesterol is an integral part of host membranes, it is likely that perturbations in cholesterol homeostasis are required for viral envelopment and egress. Cholesterol depletion from the host cell plasma membrane using methyl-β-cyclodextrin broadly inhibits a diverse array of viruses (43–52) and methyl-β-cyclodextrin treatment prevents KSHV egress by blocking secondary envelopment at the TGN (53). In addition, multiple KSHV-encoded microRNAs expressed during latency target and repress multiple steps in the cholesterol biosynthetic pathway (54, 55) However, exogenous regulation of intracellular cholesterol levels by lovastatin or 25-hydroxycholesterol lead to higher latently infected PEL cell death or restricted *de novo* KSHV infection (56, 57). These studies clearly implicate cholesterol regulation in various aspects of the KSHV life cycle, but more study is required to understand the precise viral components that interact with cholesterol during KSHV lytic replication.

One of the main regulators of cellular cholesterol homeostasis is liver X receptor alpha (LXRα). LXRα is a nuclear transcription factor responsible for the transcription of genes containing LXR response elements (LXREs), including cholesterol storage and efflux genes (58). One of the classical LXR response genes is ATP binding cassette subfamily A member 1 (ABCA1), which contains 12 transmembrane helices folded into two distinct transmembrane domains and is localized to the plasma membrane to control cholesterol export to nascent high-density lipoprotein particles (59–63). LXRα is activated under cholesterol-rich conditions by oxidized or enzymatically-derived products of cholesterol known as oxysterols (64). When added exogenously, oxysterols stimulate cholesterol efflux and alter cellular signalling pathways, where many oxysterols have been demonstrated to suppress enveloped and non-enveloped virus infection (65). In this study we used 22(*R*)-hydroxycholesterol (22OH; **Fig. 1A**), a potent oxysterol agonist of LXRα and LXRβ (66, 67), to assess the impact of cholesterol efflux on KSHV latent infection and lytic reactivation. This work, for the first time, characterizes an inhibitory role for cholesterol dysregulation on KSHV and HSV nuclear egress and illustrates the broad effects of cholesterol efflux on the host proteome.

**Figure 1.**
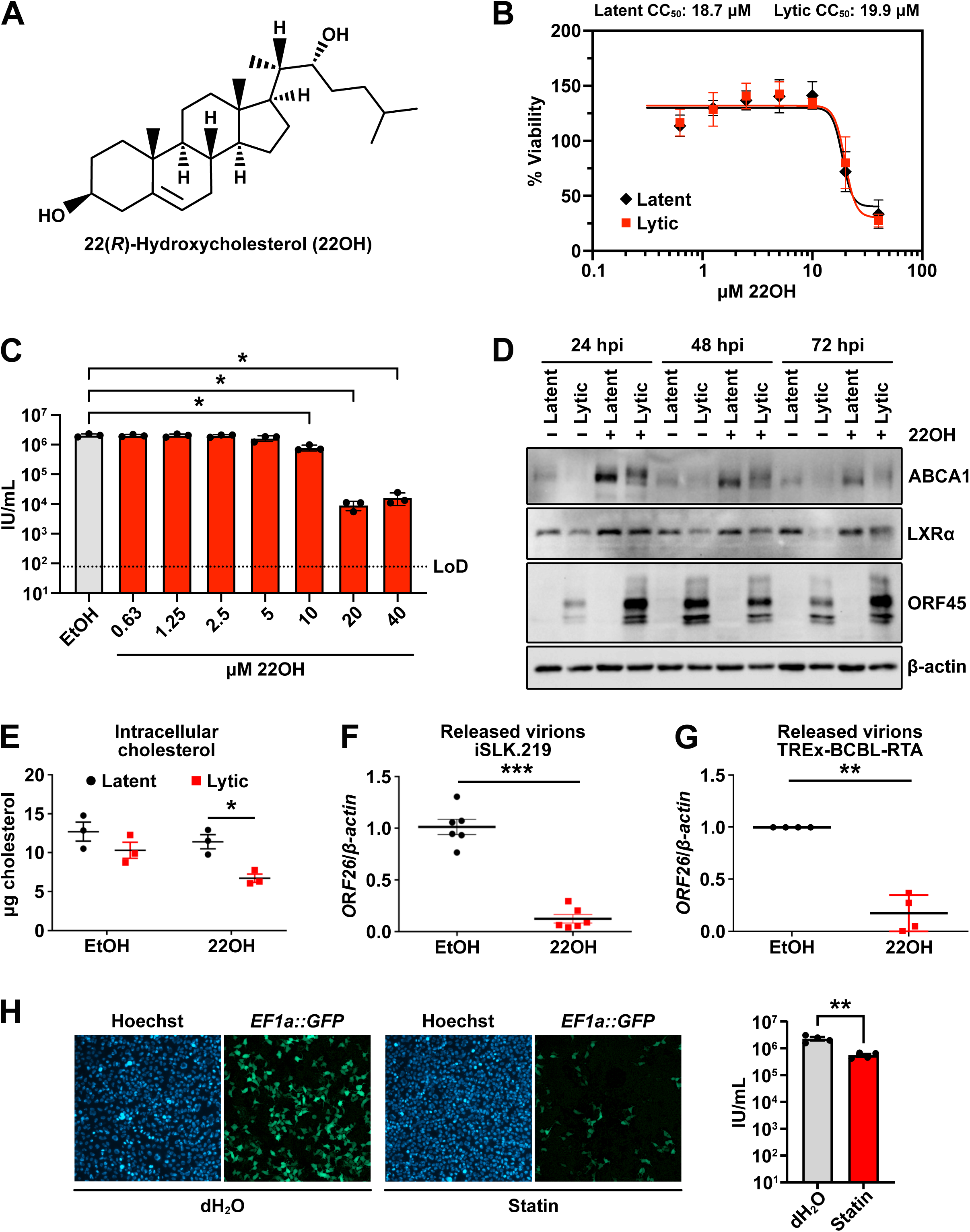
Intracellular cholesterol depletion reduces KSHV production. **(A)** Chemical structure of 22(*R*)-hydroxycholesterol (22OH). **(B)** Cell viability following 22OH treatment. iSLK.219 cells were treated with vehicle control or doxycycline (dox) to induce lytic replication, and with vehicle control or a range of concentrations of 22OH from 0.63 µM to 40 µM. Cell viability was measured by CCK-8 assay at 48h post-treatment; n = 3. **(C)** Dose-dependent inhibition of KSHV by 22OH. Cell supernatants were collected at 96h post-treatment from a dilution series experiment similar to panel A and titered by TCID50 assay; n = 3, \**p* < 0.05. **(D)** 22OH upregulates ABCA1 in iSLK.219 cells. iSLK.219 cells were treated with vehicle control or 9 µM 22OH to induce cholesterol efflux and left untreated (Latent) or treated with doxycycline to induce reactivation (Lytic). Protein lysates were prepared at the indicated times post-reactivation for SDS-PAGE and immunoblotting for ABCA1, LXRα, KSHV ORF45, or β-actin. **(E)** 22OH decreases total cellular cholesterol in lytic cells. iSLK.219 cells were treated with vehicle control (latent, black symbols) or dox to induce reactivation from latency (lytic, red symbols), as well as 9 µM 22OH or vehicle control for 24 h prior to harvest in phosphate-buffered saline with 0.1% SDS (w/v) and processed with an Amplex Red cholesterol assay to measure total esterified cellular cholesterol; n=3, \**p* < 0.02. **(F,G)** 22OH inhibits release of KSHV particles from iSLK.219 and TREx-BCBL-RTA cells. **(F)** iSLK.219 cells were treated with dox to induce lytic reactivation, and 9 µM 22OH to stimulate cholesterol efflux. Cell supernatants were harvested at 96h post-treatment and capsid-protected genomes were measured by qPCR; n = 6, *** *p* < 0.005. **(G)** TREx-BCBL-RTA cells were treated with dox to induce lytic reactivation, and 9 µM 22OH to stimulate cholesterol efflux. Cell supernatants were harvested at 48h post-treatment and capsid-protected genomes were measured by qPCR; n = 4, **** *p* < 0.001. **(H)** Statin treatment inhibits KSHV production. Supernatants of iSLK.219 cells treated with type II water (dH_2_O) or 50 nM cerivastatin were used to infect 293A cells and infected cells were imaged 24 h later to visualize Hoechst 33342 (DNA, blue) and green fluorescent protein (GFP) expression (viral reporter, green) (left panel). KSHV titres calculated from automated image analysis are plotted as the mean ± standard error of the mean for each treatment from four biological replicates. **p < 0.01 (right panel).

## RESULTS

### Reduced intracellular cholesterol inhibits KSHV production

To investigate the effect of cholesterol efflux on KSHV infection, latently KSHV infected iSLK.219 cells were mock treated or treated with doxycycline to trigger lytic replication and concurrently treated with a range of 22OH doses from 40 μM down to 0.64 μM via 2-fold serial dilution. Cell viability was measured at 48 h post-treatment and demonstrated equivalent effects of 22OH on cell viability during latent (CC_50_ = 18.73 μM) and lytic infection (CC_50_ = 19.87 μM) (**Fig. 1B**). Titration of cell supernatants harvested from iSLK.219 cells at 96 h post-treatment demonstrated a dose-dependent inhibition in release of infectious virions with 22OH treatment (IC_50_ = 8.56 μM; **Fig. 1C**). Using a sub-cytotoxic dose of 22OH (9 µM), we observed that treatment of latently KSHV infected iSLK.219 epithelial cells with 22OH caused a marked increase in protein production from LXRα target gene ATP binding cassette subfamily A member 1 (ABCA1) (**Fig. 1D**), indicating that LXRα can respond as expected to an oxysterol agonist in these cells. Doxycycline-mediated reactivation from latency, reflected by the accumulation of the early KSHV protein ORF45, correlated with decreases in ABCA1 and LXRα proteins, consistent with host shutoff; however, these decreases were moderated by 22OH treatment (**Fig. 1D**). We also observed that 22OH treatment during KSHV lytic replication altered the electrophoretic mobility of ABCA1 (**Fig. 1D**), which is consistent with known regulation of ABCA1 by phosphorylation, palmitoylation, glycosylation, and ubiquitination (68–71). Consistent with our observation of upregulation of ABCA1 by 22OH, we also observed significantly reduced total cellular cholesterol (esterified and unesterified) in lytically infected cells treated with 22OH and no significant effect on latent iSLK.219 cells (**Fig. 1E**). Thus, in KSHV infected cells, the LXRα-ABCA1 pathway is intact and 22OH can stimulate cholesterol efflux.

Next, we used a qPCR assay to quantify release of capsid-protected genomes (a measure of viral particles rather than infectivity) from dox-treated iSLK.219 cells and TREx-BCBL-RTA cells, which revealed a ∼10-fold drop in viral particles in cell supernatants in both infection models (**Figs. 1F** and **1G**). This suggests that 22OH treatment inhibits the production of viral particles by KSHV infected cells but does not pinpoint which post-entry step in lytic replication is sensitive to the drug.

To ensure that the effects of 22OH treatment are not related to off-target effects, we assessed the impact of treating iSLK.219 cells with cerivastatin; a potent 3-hydroxy-3-methylglutaryl-coenzyme A reductase (HMGCR) inhibitor that prevents *de novo* cholesterol synthesis of new cholesterol leading to a net reduction in total intracellular cholesterol levels (72–74). After performing a 96 h infection of iSLK.219 cells in the presence or absence of cerivastatin (statin), the virus containing supernatants were collected, used to infect naïve 293A cells, and GFP-positive cells were counted relative to Hoechst staining (DNA, nuclei) to assess the number of infectious KSHV particles present. Cerivastatin treatment resulted in a 4-fold reduction in infectivity from the supernatants compared to the water control supernatants (**Fig. 1H**). These results demonstrate that reduced cellular cholesterol levels through reduced cholesterol production can also lead to significant declines in the production of infectious virions.

### 22OH increases reactivation from latency but diminishes viral protein accumulation at late times post-reactivation

iSLK.219 cells are latently infected with a recombinant KSHV that constitutively expresses green fluorescent protein (GFP) from an elongation factor 1α (*EF1α*) promoter; the addition of doxycycline increases the synthesis of the replication and transcription activator (RTA) lytic switch protein, thereby triggering reactivation from latency and expression of a red fluorescent protein (dsRed) reporter gene driven by the KSHV *PAN* promoter. Treatment of latently infected cells with 22OH alone did not elicit reactivation from latency (**Figs. 2A-B**). By contrast, co-administration of 22OH with doxycycline caused a significant increase in red fluorescent cells in the monolayer at 24 and 48 h post-reactivation compared to doxycycline treatment alone as visualized by immunofluorescence microscopy and flow cytometry (**Figs. 2A-B**). This suggested that 22OH could potentiate reactivation from latency in the presence of sufficient levels of RTA. This was consistent with our observation by immunofluorescence of increased numbers of cells expressing the early lytic protein ORF57 following treatment with doxycycline and 22OH at 24 h post-reactivation compared to doxycycline treatment alone (**Fig. 2C**). Viral genome replication in this cell population was significantly increased in 22OH treated cells at 48 h post-reactivation (**Fig. 2D**), consistent with increased frequency of reactivation from latency and unimpeded progression to the viral genome replication step. By 72 h post-reactivation, no significant differences in viral genome replication remained between the mock-treated and 22OH-treated cells (**Fig. 2D**). The herpesvirus DNA polymerase inhibitor phosphonoacetic acid (PAA) served as a positive control for inhibition of viral DNA replication in this assay (**Fig. 2D**).

**Figure 2.**
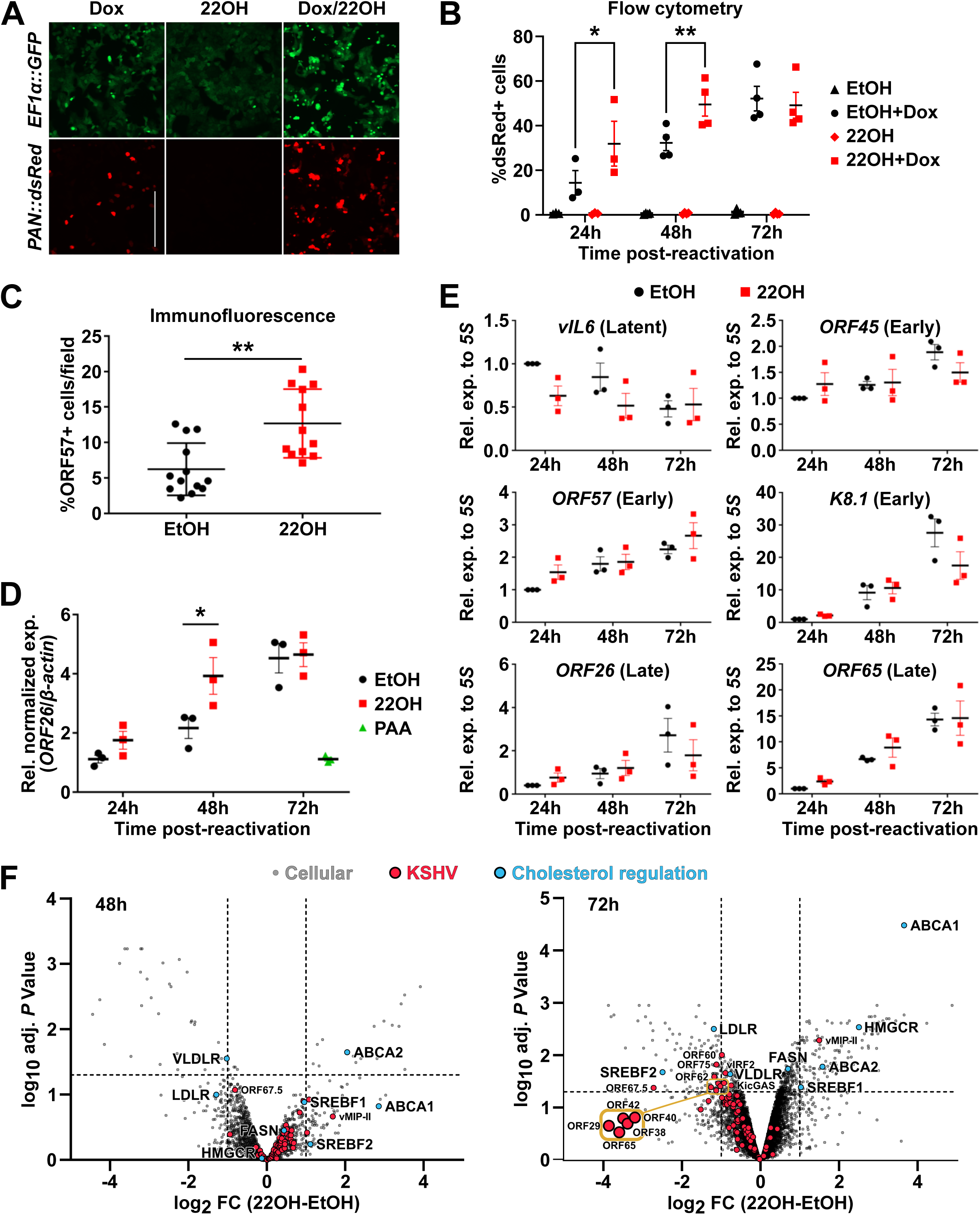
Cholesterol efflux increases KSHV reactivation from latency but diminishes viral protein accumulation at late times post-reactivation. **(A-C)** 22OH increases reactivation from latency. iSLK.219 cells were reactivated with 1 µg/mL doxycycline (Dox) and treated with 10 µM 22OH. Live cell fluorescence microscopy was conducted 24 h post-reactivation **(A)** or flow cytometry was performed at 24, 48, and 72 h post-reactivation; n = 3-4, * *p* < 0.05, ** *p* < 0.01 **(B)**; green fluorescence (GFP) reports latent infection and red fluorescence (dsRed) reports early lytic replication. **(C)** iSLK.219 cells were fixed 24 h post-reactivation and immunostained with anti-ORF57 antibody and DAPI (DNA/nuclei) and expressed as %ORF57-positive cells; n=3, *p* = 0.0013. **(D)** 22OH does not inhibit viral genome replication. DNA was harvested from iSLK.219 cells at times indicated post-reactivation and intracellular KSHV genomes were enumerated by qPCR amplification of *ORF26* normalized to *β-actin*. In parallel, cells were treated with phosphonoacetic acid (PAA) to block genome replication. A 2-way ANOVA was performed to determine significance n = 3, \**p* = 0.02. **(E)** 22OH does not affect accumulation of representative early and late viral mRNAs. RNA was harvested from iSLK.219 cells at times indicated post-reactivation; RT-qPCR was performed on viral genes indicated. **(F)** 22OH decreases accumulation of viral proteins at late times post-reactivation. Whole proteomes from iSLK.219 cells at 48 h or 72 h post-reactivation with or without 10 µM 22OH treatment are shown. Viral proteins are in red, with viral nuclear egress complex (NEC) proteins in gold, and proteins responsive to 22OH and involved in cholesterol regulation are shown in blue. Host proteins are indicated in grey. These data are plotted with the log_2_ fold-change (FC) of 22OH-treated relative to ethanol (EtOH)-treated on the x-axis and the log_10_ adjusted (adj.) *P* value on the y-axis.

To pinpoint the stage of herpesvirus lytic replication affected by treatment with 22OH, we analysed viral mRNA and protein accumulation. iSLK.219 cells were reactivated in the presence of vehicle control or 22OH and cell lysates were harvested at 0, 24, 48 and 72 h post-reactivation. We observed that 22OH treatment did not significantly affect accumulation of representative viral latent mRNAs (*vIL-6*), early mRNAs (*ORF45* and *ORF57*), delayed-early mRNAs (*K8.1*), or late mRNAs (*ORF26* and *ORF65*) ((75); **Fig. 2E**), consistent with our previous observations of unimpeded viral genome replication in treated cells (**Fig. 2D**). Recognizing the limitations of interpreting effects of 22OH on viral protein accumulation with a limited number of available antibodies, we harvested lysates from vehicle-treated or 22OH-treated latent iSLK.219 cells at 48 h and 72 h post-reactivation and processed them for label-free proteomics. The effects of 22OH on LXRα targets was appreciable at 48 h post-reactivation and more pronounced by 72 h post-reactivation, with significantly altered expression of multiple proteins involved in regulating cholesterol homeostasis (blue dots), including: SREBF2, LDLR, VLDLR, FASN, HMGCR, as well as ABCA1 and ABCA2 (**Fig 2F**). The net effect of 22OH treatment reducing cellular cholesterol results from both the downregulation of cholesterol uptake proteins (i.e. LDLR and VLDLR) and the upregulation of cholesterol efflux proteins (i.e. ABC proteins) (76). We also observed upregulation of ABCA1 by western blot (**Fig. 1D**). We observed modest 22OH-induced increases in the accumulation of most viral proteins (red dots) at 48 h (**Fig. 2F**, left panel), consistent with our observations of increased frequency of lytic reactivation in these cells (**Figs. 2A-C**). Interestingly, despite the clear stimulation of KSHV reactivation in the presence of 22OH treatment, levels of the RTA protein are relatively unchanged at 48 h (**Table S1**), making it unlikely that enhanced reactivation is solely due to increased production of RTA. By 72 h post-reactivation, ABCA1 was the sixth most upregulated host protein (**Fig. 2F**, right panel and **Table S2**). At this late time point, viral protein levels also showed broad decline in the 22OH-treated cells compared to mock-treated counterparts, which is consistent with the observed reduction in the release of infectious progeny (**Figs. 1F-G**). The viral proteins most negatively affected by 22OH treatment by 72 h post-reactivation were (ordered by decreasing log_2_ FC): ORF67.5 (terminase complex (77)), ORF29 (terminase complex (78)), ORF62 (capsid protein (79)), ORF65 (small capsid protein (79)), ORF75 (vFGARAT (80)), ORF42 (tegument protein (81)), ORF38 (tegument protein (82)), ORF60 (small vRNR (80)), ORF40 (helicase (80)), K11 (vIRF2 (83)), and ORF52 (KicGAS (84)) (**Fig. 2F**, right panel and **Table S2**; *p* values < 0.05). Interestingly, the viral cytokine vMIP-II (encoded by K4 (85)) was significantly upregulated following 22OH treatment (**Fig. 2F**, right panel; log_2_ FC: 1.50, *p* value: 0.005). And of the most highly affected viral proteins at 72 h, both ORF67.5 and vMIP-II were also affected by 22OH at 48 h post-reactivation; however, these effects were not significant at this earlier timepoint (**Fig. 2F**, left panel). Altogether, the effects of 22OH on viral protein expression are broad and affect multiple proteins required for mature virion formation and immune regulation.

### 22OH treatment elicits a non-conventional innate immune response

Previously, the antiviral activity of a related oxysterol, 25-hydroxycholesterol (25OH), during KSHV *de novo* infection of human umbilical vein endothelial cells (HUVECs) was assessed using RNA-seq analysis and demonstrated that, broadly, interferon stimulated genes (ISGs) and inflammatory genes were upregulated at the mRNA level (57). This led us to postulate that 22OH could have similar effects on innate immune responses in iSLK.219 cells. In comparing our proteomics dataset to the previous RNA-seq data, we observed that the proteins MAFF (48 h), ATF3 (72 h) and FOSL1 (72 h) were upregulated by 22OH just as they were transcriptionally induced by 25OH (**Tables S1 and S2** (57)). Our proteomics datasets had sufficient depth with 8985 (48 h) and 8888 (72 h) identified peptides (including KSHV proteins) to allow us to evaluate global changes to the proteome following 22OH treatment of infected iSLK.219 cells using Gene Set Enrichment Analysis (GSEA) (86, 87). GSEA of our proteomics data did not demonstrate an activation of a classical inflammatory response as observed with 25OH treatment (57); instead, we observed more enrichment in the downregulation of proteins involved in the type I interferon (IFN) and inflammatory responses compared to upregulation (**Fig. 3A**). We did not detect type I IFNs or other antiviral cytokines in our proteomics dataset (**Tables S1 and S2**). These differences in responses to 22OH and 25OH could be due to their differing affinities for LXRs (66, 67). Broadly, 22OH treatment also had significant impacts on cellular signalling pathway proteins, with notable enrichment (upregulation) in proteins involved in regulating the cell cycle (**Fig. 3A**).

**Figure 3:**
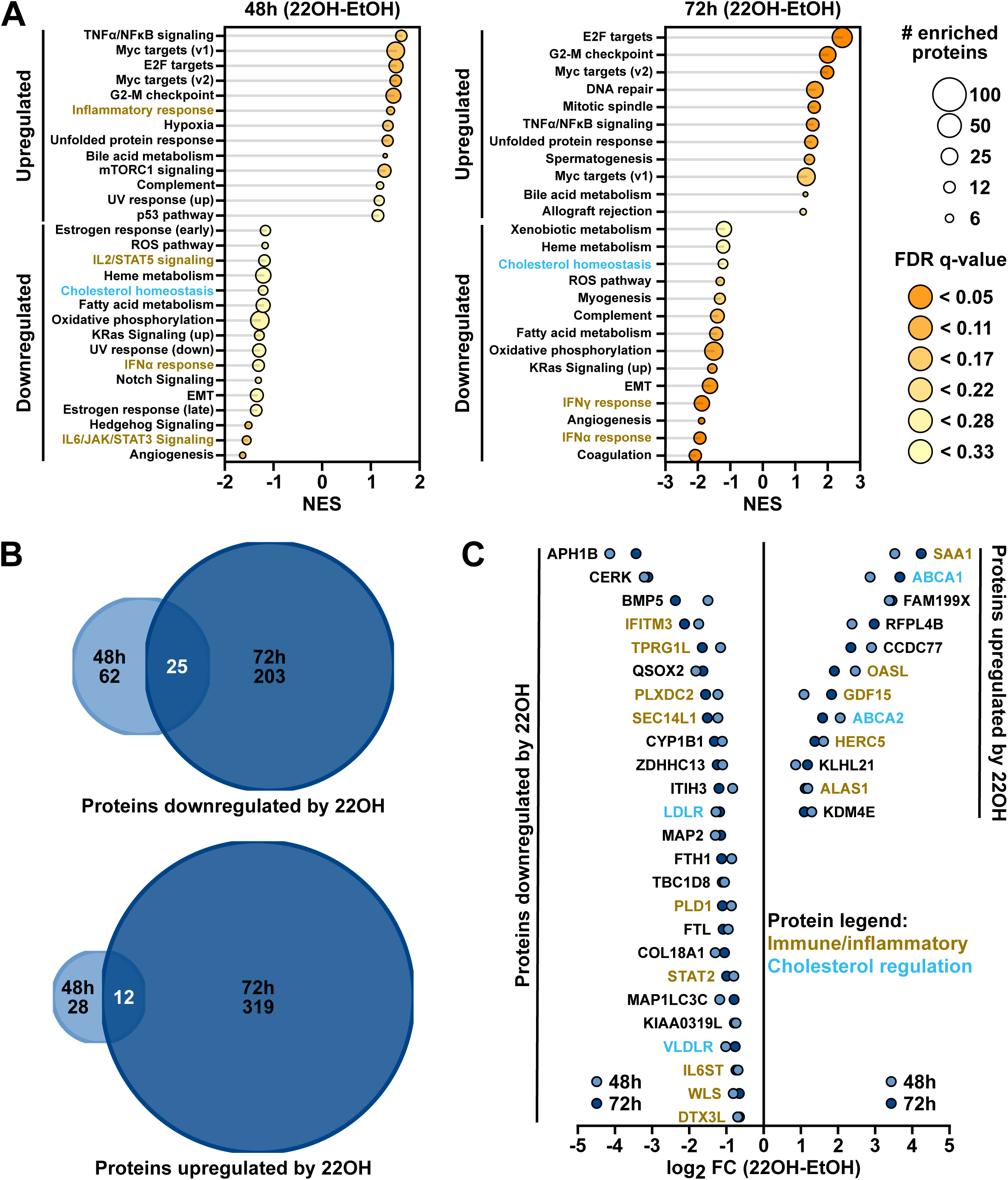
22OH treatment elicits a non-conventional innate immune response. **(A)** Gene Set Enrichment Analysis (GSEA) was performed using ranked lists ordered by log_2_ fold-change (log_2_ FC) of the 48 h and 72 h proteomic datasets shown in Fig. 2F. All classifications with false discovery rate (FDR) q-values of < 0.33 are shown and plotted against the normalized enrichment score (NES). Circle size indicates the number of proteins enriched in each classification and colour-coding illustrates the FDR q-value for each classification. Immune and inflammatory classifications are highlighted with gold font and cholesterol homeostasis is highlighted in light blue font. **(B)** Area-proportional Venn diagrams illustrating the number of significantly downregulated (top) or upregulated (bottom) cellular proteins from each dataset at either 48 h (light blue) or 72 h (dark blue) based on spectra count-adjusted adjusted *P*-values < 0.05 and log2 FC values of < −0.6 (downregulated) or > 0.6 (upregulated). The number of cellular proteins downregulated or upregulated at both time points is shown in white font. **(C)** The shared downregulated and upregulated cellular proteins from the 48 h (light blue) and 72 h (dark blue) proteomic datasets are shown plotted with their calculated log_2_ FC. Cellular proteins known to participate in immune or inflammatory responses are in gold font and those that participate in cholesterol regulation are shown in light blue font.

For each proteomics dataset, which comprised three independent experiments per timepoint, we identified our most significant downregulated and upregulated proteins based on spectra count-adjusted adjusted *P*-values < 0.05 and sorted these by log2 FC values with conservative cutoffs at −0.6 (downregulated) and > 0.6 (upregulated). Comparing the lists of upregulated or downregulated proteins between the two timepoints, we observed 25 proteins consistently downregulated by 22OH and 12 proteins consistently upregulated by 22OH (**Figs. 3B** and **3C**). Notably, there was reduced expression of multiple proteins either directly or indirectly involved in innate immune responses, including: IFITM3, TPRG1L, PLXDC2, SEC14L1, PLD1, STAT2, IL6ST, WLS, and DTX3L (**Fig. 3C** and **Table S3**). However, some innate immune regulatory proteins and proteins involved in inflammatory responses were upregulated, including: SAA1, OASL, GDF15, HERC5, and ALAS1 (**Fig. 3C** and **Table S3**). While we do not observe the hallmarks of a classical innate immune response due to 22OH treatment in reactivated iSLK.219 cells, there are characteristics of an atypical antiviral response being induced.

### Cholesterol efflux affects KSHV capsid assembly and packaging

With our observations from our proteomics data that identified potential inhibitors of KSHV replication (**Fig. 3C**), as well as significant reductions in KSHV proteins involved in virion assembly and genome packaging (**Fig. 2F**), we next wanted to assess the impact of 22OH treatment on KSHV virion production. Our previous experiments demonstrated diminished release of viral particles from two different KSHV model systems (**Figs. 1F-G**), but these data did not indicate the point in the viral life cycle where the defect occurs.

To investigate potential defects in KSHV assembly and egress, we harvested and processed infected cells for negative staining and transmission electron microscopy (TEM) analysis. During herpesvirus replication, an early process in the generation of new viral progeny involves the generation of protein capsids needed for the encapsulation of viral genomes. In a cell undergoing lytic replication, there are three capsid species that predominate: A-capsids (empty, no genome or inner scaffold), B-capsids (contain inner scaffold only), and C-capsids (mature structure containing inner scaffold and viral genome) (79). A- and B-capsids are considered dead-end byproducts of capsid assembly and only C-capsids are actively recruited for subsequent nuclear egress. Our TEM experiments were performed with three independent biological replicates and >900 capsids were enumerated from multiple sections per replicate. As expected, no viral capsids were observed in latently infected iSLK.219 cells treated with 22OH or vehicle control for 24 h (**Fig. 4A**). At 48 h post-reactivation, KSHV capsids accumulated in nuclei of vehicle-treated cells, most of which were C-capsids (51%, blue), followed by B-capsids (35%, red), and A-capsids (14%, yellow; **Figs. 4A-B**); 22OH-treated cells displayed a slightly different distribution of capsid types with a decline in the proportion of C-capsids (35%; **Fig. 4B**). At 72 h post-reactivation, C-capsids predominated once again in vehicle-treated cells (**Fig. 4B**). However, at this time point, C-capsid numbers were reduced in 22OH-treated cells, with A-capsids emerging as the predominant capsid type. Thus, 22OH treatment altered the proportions of KSHV capsid types at later stages of infection with a shift towards immature or defective A/B-capsids (79) in the nuclei of 22OH treated cells.

**Figure 4:**
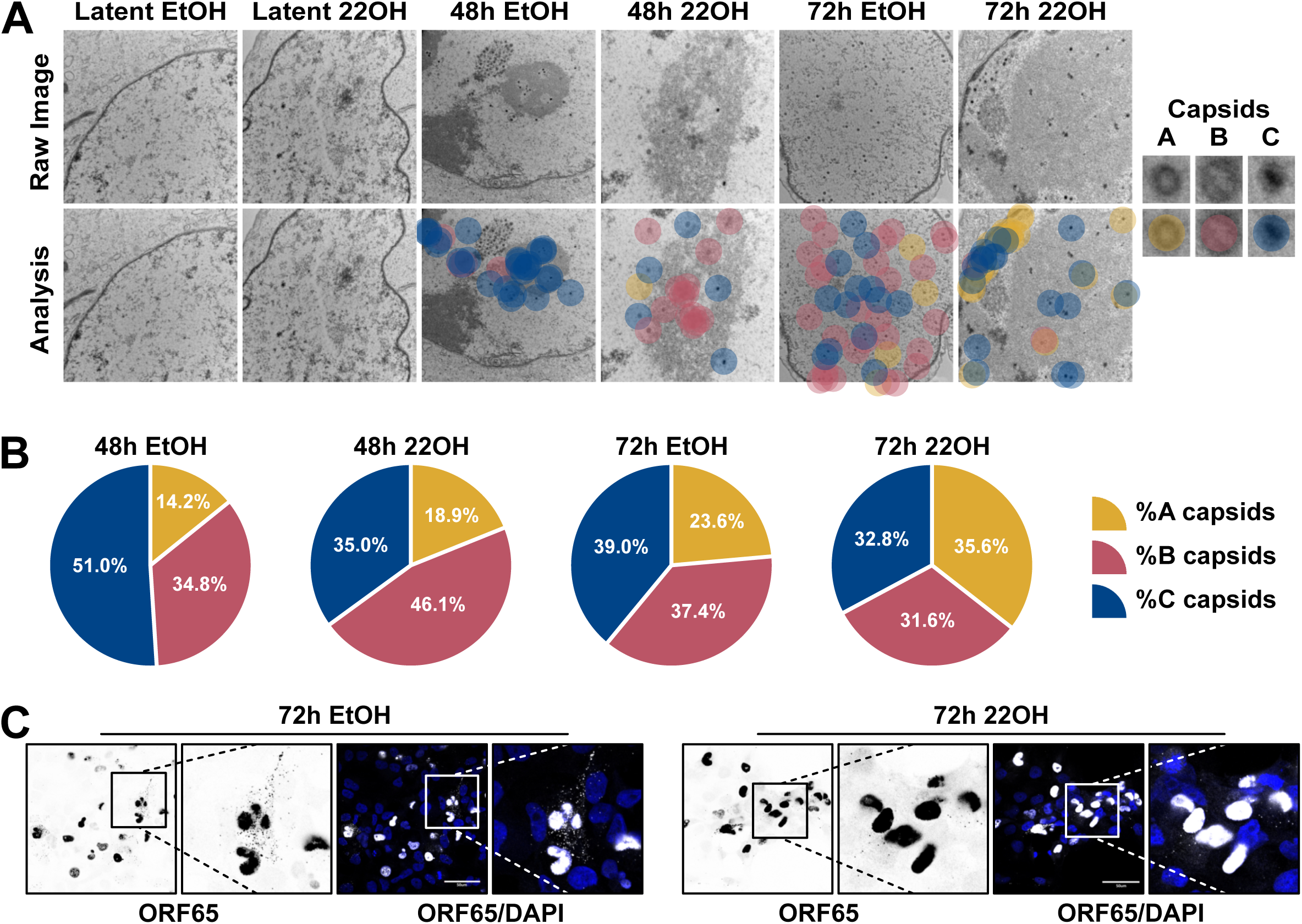
LXRα activation inhibits KSHV nuclear egress. KSHV nuclear egress was monitored using iSLK-BAC16 cells reactivated from latency with 1 µg/mL doxycycline and 1 mM sodium butyrate in the presence of 22OH (10 µM) or vehicle (ethanol, EtOH) control. **(A)** iSLK-BAC16 cells not treated with doxycycline (latent, 48 h) or those treated with doxycycline were harvested at 48 h and 72 h post-reactivation and fixed and processed for TEM. A, B, and C capsids were enumerated in all images as indicated by the yellow (A-capsids), red (B-capsids), and blue (C-capsids) notations in representative images from three independent biological replicates. Examples of each capsid type are shown to the far-right. **(B)** Capsids were quantified from all images and replicates and were plotted as percentages of the total number of enumerated capsids for each treatment condition. Total capsids counted: 48 h EtOH, 719; 48 h 22OH, 401; 72 h EtOH, 1126; 72 h 22OH, 747. **(C)** iSLK.219 cells seeded onto coverglass were fixed at 72 h post-reactivation and stained with anti-ORF65 (late lytic, small capsid protein) antibody and DAPI (DNA/nuclei). The inset panels are shown with increased magnification to the right to highlight cytoplasmic ORF65 punctae.

Quantitative surveys of cellular compartments by conventional TEM are limited by the lack of information in the Z-plane. To overcome this limitation and investigate KSHV nuclear egress and capsid accumulation in the cytoplasm, we immunostained reactivated iSLK.219 cells with an antibody against the small capsid protein (ORF65) to mark the position of cytoplasmic capsids and presented these images as maximum intensity projections to visualize the entire cytoplasmic volume. We readily detected the accumulation of ORF65-containing KSHV capsids in both the nucleus (strong, condensed staining) and cytoplasm (lower intensity punctate staining) of vehicle-treated infected cells (**Fig. 4C**). In 22OH-treated cells, the nuclear ORF65 signal remained strong; however, the quantity of cytoplasmic ORF65 puncta was greatly reduced in 22OH-treated cultures (**Fig. 4C**). With 22OH treatment, by 72 h post-reactivation ORF65 protein levels were reduced to 45% (log_2_ FC: −1.12, *p* value: 0.02) or those in vehicle treated cells (**Fig. 2F** and **Table S2**). As mentioned above, components of the viral terminase complex (ORF29 and ORF67.5) were also reduced following 22OH treatment (**Fig. 2F** and **Table S2**). Together, these observations suggest that stimulating cholesterol efflux impinges on KSHV nuclear egress by disrupting capsid assembly or packaging.

### 22OH inhibits herpes simplex virus infection and alters viral nuclear egress complex localization

To determine whether the antiviral effects of cholesterol efflux are conserved amongst distantly related herpesviruses, we also tested 22OH on herpes simplex virus type 1 (HSV-1) and type 2 (HSV-2). Using a plaque reduction assay, we observed that 22OH significantly inhibited both plaque number and plaque size across a range of concentrations from 10 μM to 0.1 μM with HSV-1 infection (**Figs. 5A-B**). Similar results were obtained in plaque reduction assays with HSV-2; however, the only dose of 22OH to yield significant reductions in plaque number and size was 10 µM (**Figs. 5C-D**). Similar to our earlier results with KSHV, the observed reduction in HSV replication did not correlate with a reduction in viral protein expression at early times post-infection (**Fig. 5E**). We investigated the expression of HSV-1 immediate-early (ICP27), early (pUS3), and late (gC) proteins with and without 22OH treatment and observed no appreciable changes in viral protein accumulation, unlike when viral DNA replication was blocked with phosphonoacetic acid (PAA) (**Fig. 5E**).

**Figure 5:**
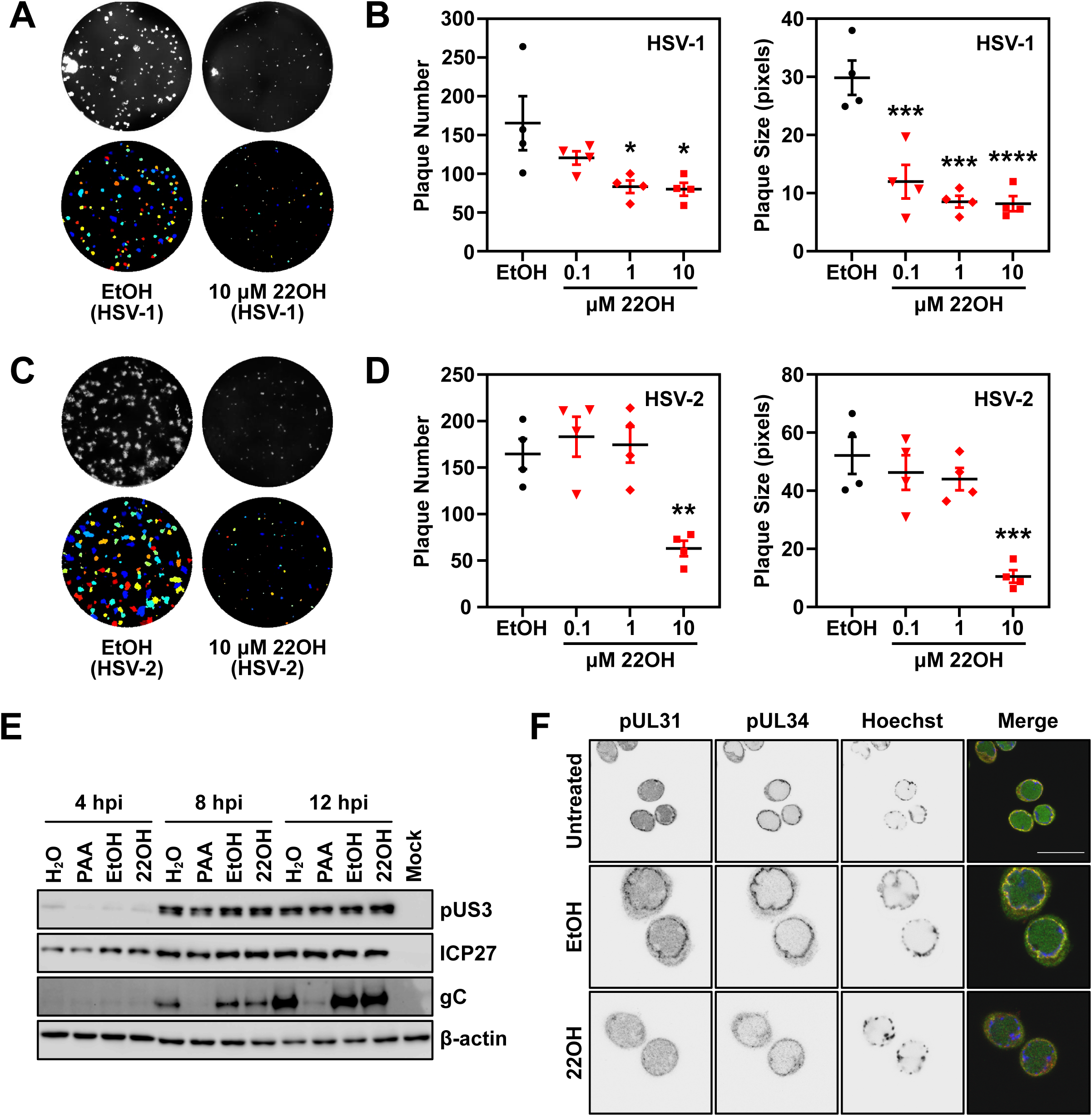
LXRα activation impairs HSV-1 and HSV-2 replication and disrupts nuclear egress complex localization to the nuclear lamina. Representative images of HSV-1 **(A)** or HSV-2 **(C)** plaque reduction assays in HeLa cells are shown with the raw images on top (black and white) and the plaque identification from CellProfiler shown on the bottom (colour). CellProfiler quantitation of the results of four independent plaque reduction assays for HSV-1 **(B)** and HSV-2 **(D)** are shown. Data were analyzed using one-way ANOVA (Tukey). \**p* < 0.05, \*\**p* < 0.01, \*\*\**p* < 0.001, \*\*\*\**p* < 0.0001. **(E)** Protein lysates were prepared from HSV-1 infected HeLa cells at the indicated hours post-infection (hpi) for SDS-PAGE and immunoblotting for HSV-1 proteins pUS3, infected cell protein 27 (ICP27), and glycoprotein C (gC) or β-actin. **(F)** Immunofluorescence microscopy of HSV-2 infected HeLa cells at 24 hpi following staining with antibodies against the nuclear egress complex proteins pUL31 and pUL34 and Hoechst 33342 (DNA). Abbreviations: 22OH, 22-hydroxycholesterol; EtOH, ethanol; H_2_O, water; PAA, phosphonoacetic acid.

Because 22OH also inhibited HSV replication in HeLa cells and we previously observed defects in nuclear egress with KSHV, we next investigated the effects of 22OH on HSV-2 nuclear egress. All herpesviruses express orthologs of HSV pUL31 and pUL34 that constitute the viral nuclear egress complex (NEC) (88). These two proteins are recruited to nuclear membranes to facilitate the egress of assembled nuclear capsids into the cytosol. When we examined the localization of pUL31 and pUL34 in HSV-2 infected cells, we observed dramatically reduced recruitment of the nuclear egress complex (NEC) proteins pUL31 and pUL34 to the nuclear periphery in 22OH treated cells (**Fig. 5F**). In our proteomics with KSHV, only the level of the pUL31 ortholog (ORF69; log_2_ FC: −0.59, *p* value: 0.03) was significantly reduced following 22OH treatment (**Table S2**). These data along with our TEM data from KHSV infected cells suggest a common mechanism for 22OH-mediated disruption of nuclear egress for divergent herpesviruses.

## DISCUSSION

Cholesterol is incorporated into herpesvirus envelopes and its presence in viral envelopes and target host membranes is required to support membrane fusion during viral entry (33, 42). By contrast, relatively little is known about how cholesterol affects post-entry processes. Here, using infection models that bypass the entry step, we demonstrated that drugs that deplete intracellular cholesterol, either by inhibiting cholesterol biosynthesis (cerivastatin) or promoting cholesterol efflux (22OH), inhibit the production of infectious KSHV virions (**Fig. 1**). In further investigating the effects of 22OH treatment, we determined that this inhibition occurs after viral genome replication and features reduced accumulation of viral proteins at 72 h post-reactivation, increased accumulation of A-capsids and B-capsids in the nucleus, and a failure of capsids to access the cytoplasm (**Figs. 2F** and **3**). Together, these findings indicate that disrupting cholesterol homeostasis interferes with KSHV nuclear egress, although precise mechanisms remain unknown. Certainly, reduced accumulation of viral proteins in late stages of replication could interfere with efficient capsid assembly, maturation, and/or genome encapsidation (see those highlighted in **Fig. 2F**), which would support our observation of a shift towards more immature/empty capsids in the nuclei of 22OH-treated cells. Cholesterol plays key roles in regulating the fluidity of cellular membranes and transmembrane protein motility, and the distribution of intracellular cholesterol has a profound impact on protein trafficking in the endomembrane system (89–91). Thus, depletion of intracellular cholesterol by stimulating efflux could interfere with the trafficking and fate of viral proteins involved in capsid assembly and nuclear egress.

The best described mechanism for KSHV lytic reactivation is endoplasmic reticulum (ER) stress-induced Inositol-Requiring Enzyme 1 (IRE1) activation and generation of the X-box-binding protein 1 spliced (XBP1s) transcription factor that transactivates the RTA lytic switch transcription factor (92, 93). Lipid regulation of the unfolded protein response (UPR) is conserved from yeast to humans; IRE1, and XBP1s (originally named IRE2) were first discovered in yeast as required for growth in the absence of inositol, a building block for phospholipids (94, 95). Our observation that 22OH stimulates reactivation from latency (**Figs. 2A-C**) leads us to speculate that treatment with oxysterols could trigger viral reactivation via lipid-stress mediated UPR activation, which directs the clustering and oligomerization of UPR sensors IRE1 and PKR-like endoplasmic reticulum kinase (PERK) (96–99). This could be directly tested by monitoring activation of IRE1 and PERK following treatment with 22OH or other oxysterols, both in latent KSHV infection models, and models of other herpesviruses like Epstein-Barr virus (EBV) which has a similar mode of ER stress-responsive lytic reactivation (100). As mentioned earlier, the understanding that cellular metabolism is dramatically altered during KSHV latency compared to uninfected cells (25) and that these changes may prime these cells for reactivation (26) could be central to our observations of enhanced reactivation in cells with low cholesterol (**Figs. 2A-C**).

Following genome encapsidation in the nucleus, herpesvirus capsids are recruited to the INM for primary envelopment. This process is facilitated by the NEC, comprised of two highly conserved viral proteins that accumulate at the INM and form a hexagonal lattice that induces membrane curvature to support capsid budding (88). Seizing an opportunity to study the best characterized NEC proteins from alphaherpesviruses, we demonstrated that 22OH treatment inhibited HSV-1 and HSV-2 replication and spread in a dose-dependent manner in plaque reduction assays, and immunostaining HSV-2 infected cells with antibodies directed against the pUL31 and pUL34 NEC proteins showed that 22OH inhibited their recruitment to the nuclear periphery (**Fig. 5**). KSHV ORF69 is known to remodel nuclear membranes in participation with ORF67 during primary envelopment (17) and our proteomics data show a reduction in ORF69 levels at late times post-reactivation (**Fig. 2F**, right panel). These observations of effects on components of HSV-2 and KSHV NECs provide additional support for a cholesterol efflux-driven herpesvirus nuclear egress defect at the INM.

We recently demonstrated that in addition to budding at the peripheral nuclear membrane, KSHV capsids can also bud into dynamic INM infoldings known as the type-I nucleoplasmic reticulum (NR) (101, in press). These Type-I NR structures co-localized with puncta containing CTP:phosphocholine cytidylyltransferase (CCTα), the enzyme that catalyzes the rate-limiting step in phosphatidylcholine synthesis that drives the *de novo* membrane biogenesis and membrane curvature required for NR expansion (102, 103). Type-I NR is lamin-poor compared to other regions of the nuclear envelope and may provide favourable sites for nucleocapsid primary envelopment. It is plausible that cholesterol may affect dynamic nuclear infolding; while the nuclear envelope is generally considered cholesterol-poor, the lamin B receptor (LBR) at the INM catalyzes cholesterol synthesis via sterol reductase activity and is vital for nuclear envelope maintenance (104–106). Because cholesterol regulates the fluidity of cellular membranes, we speculate that promoting cholesterol efflux may affect nuclear membrane properties relevant to nuclear egress, like dynamic Type I NR formation or the fluidity of the outer nuclear membrane. It is also possible that cholesterol depletion could be similarly affect membranes and proteins needed for secondary envelopment following capsid egress from the nucleus. Additional experiments looking at this stage in viral replication would be needed to address this possibility.

Likely, the effects on KSHV replication by 22OH are further compounded by the defects in viral protein accumulation and induction of antiviral proteins or the suppression of proviral factors. 22OH causes an approximate 10-fold reduction in viral particle release and viral titre (**Fig. 1**). Of the viral proteins we noted to be significantly reduced by 22OH, KSHV mutant viruses defective in the production of ORF29 (2 log_10_), ORF38 (1 log_10_), ORF42 (1.5 log_10_), ORF52 (1 log_10_), ORF62 (1-3 log_10_, mutation-dependent), ORF65 1.5 log_10_), ORF67.5 (2 log_10_), and vIRF2 (1 log_10_) all display at least a 10-fold reduction in viral titre (107, 82, 84, 81, 108, 77, 78), whereas the mutant virus for vFGARAT could not be rescued (109). Notably, viral mutants for KSHV ORF29, ORF65, ORF67.5 and a rhesus monkey rhadinovirus ORF52 mutant virus (orthologous to KSHV KicGAS) all display nuclear egress defects (107, 110, 77, 78), suggesting that the smaller but cumulative reduction in the protein levels of one or all of these proteins may be central to the 22OH-mediated nuclear egress defect.

Moreover, the constellation of host factors that are collectively dysregulated by 22OH could equally factor into the observed oxysterol-mediated antiviral effect. IFTIM3 is known to be induced by KSHV infection and can inhibit fusion by viral glycoproteins in certain cell lines (111) and is broadly antiviral (112). However, the gammaherpesviruses KSHV, EBV, and murine gammaherpesvirus 68 can use a related protein, IFITM1, to support infection (113, 114), so it is possible that a downregulation in IFITM3 could similarly have pleiotropic effects depending on the cell lines used. Another highly downregulated host protein of interest is SEC14L1. This protein acts as negative regulator of RIG-I and, in its absence, could potentially lead to enhanced RIG-I mediated antiviral responses (115). Despite being a DNA virus, KSHV is sensed by RIG-I and silencing of RIG-I leads to enhanced lytic reactivation (116, 117); a phenotype that is strikingly similar to our reactivation-without-virus-production phenotype in 22OH treated cells. This is at odds with our system which one would presume has enhanced RIG-I function due to the lack of the repressor, SEC14L1. However, an additional regulator of RIG-I, DHX58/LGP2, is also downregulated due to 22OH treatment a 72 h post-reactivation (log_2_ FC: −1.04, *p* value: 0.01), which could further alter the balance of RIG-I control in infected cells. Finally, two interesting known antiviral proteins stimulated by 22OH are OASL and HERC5 (increased 3.7 and 2.6-fold, respectively; **Table S2**). During lytic reactivation, the KSHV protein ORF20 associates with OASL to facilitate viral mRNA translation while also preventing innate immune regulation by OASL (118). We speculate that during 22OH-mediated global downregulation in viral protein accumulation and the enhancement of OASL expression, that this environment could tip the balance towards facilitating an OASL-mediated antiviral responses. As the previous experiments looking at the proviral role of OASL where performed with a predominantly lytic variant of KSHV (118), it would interesting to perform OASL knockdown experiments in iSLK.219 cells in the presence of 22OH treatment to see if this blunts or abrogates the antiviral effect. Another 22OH-regulated protein that was identified in our dataset is the ISG15 E3 ligase HERC5, which is known to negatively regulate viral replication through its modification of viral and cellular proteins (119). KSHV actively disrupts the ISGylation pathway through the interaction of vIRF1 and HERC5, where the knockdown of ISG15 or HERC5 can boost KSHV production (120). Therefore, with the significant upregulation of HERC5 due to 22OH treatment, this could overcome the inhibitory activity of vIRF1. While not discussed much in this paper, the intriguing connection between 22OH treatment and cell cycle regulation will also be interesting to explore in future studies. It is plausible that 22OH-induced dysregulation of the cell cycle, over and above changes induced by herpesvirus replication, could disrupt the nuclear lamina in a way that makes it less conducive to allow capsid egress from nuclei.

Our study highlights how the cellular response to changes in intracellular cholesterol levels can influence KSHV reactivation and the importance of maintaining these levels to support productive viral infection. Despite a reduction in intracellular cholesterol levels boosting viral reactivation, subsequent stages in the viral life cycle were impaired as the burden of cholesterol loss increased at late times post-reactivation. This did not halt viral replication but reduced the accumulation of mature nucleocapsids and reduced the release of infectious KSHV. In taking a broader perspective on 22OH-mediated inhibition, we uncovered a complex cellular response to the loss of cholesterol that both reduces viral protein accumulation and dysregulates cellular signaling networks involved in the cell cycle and immune responses. Importantly, we saw no hallmarks of a classical innate antiviral response and no observable upregulation of antiviral cytokines in the presence of 22OH; instead, we saw the induction or inhibition of a variety of putative antiviral and proviral proteins. Our future studies will aim to identify the key cellular factors that contribute to the bulk of 22OH-mediated viral inhibition to better understand how the net effect of cholesterol loss makes host cells less favourable for viral replication.

## MATERIALS AND METHODS

### Cell lines

iSLK.219 cells (a kind gift from Don Ganem; (121)) are an epithelial cell line carrying a Tet-inducible RTA and are infected with recombinant KSHV rKSHV.219 (122). iSLK-BAC16 cells (a kind gift from x) are iSLK cells infected with KSHV-BAC16 (123). Treatment of iSLK cells (harbouring KSHV 219 or BAC16) with 1 µg/mL doxycycline (Sigma-Aldrich, D9891) reactivates KSHV from latency and induces lytic replication. iSLK.219 cells were cultured in 10 µg/mL puromycin (Thermo Fisher, A1113803) to maintain recombinant KSHV episome copy number. iSLK-BAC16 cells were cultured with 400 µg/mL hygromycin B (Thermo Fisher, 10687010) and 1 µg/mL puromycin. iSLK.219, iSLK-BAC16, 293A, HEK293T, and HeLa cells were cultured in 10% CO2 in Dulbecco’s Modified Eagle Medium (DMEM; Thermo Fisher, 11965118) with 10% v/v Fetal Bovine Serum (FBS; Thermo Fisher, A31607-01) and supplemented with 100 U/mL penicillin-streptomycin and 100 µg/mL L-glutamine (Pen/Strep/Gln; Thermo Fisher, 15140122 and 25030081). Vero cells were cultured in DMEM supplemented with 5% FBS and Pen/Strep/Gln. TREx-BCBL1-RTA cells (a kind gift from Jae Jung; Nakamura H., et al. 2003), were cultured in RPMI-1640 (Thermo Fisher, 11875119) with 10% FBS, 55 µM 2-mercaptoethanol (Gibco, 21985023) and Pen/Strep/Gln.

### Cell viability assay

iSLK.219 cells were seeded into 96-well tissue culture plates in 100 µL medium per well. The following day medium was replaced with fresh medium with 1 µg/mL doxycycline (lytic) or without doxycycline (latent) and 22(*R*)-hydroxycholesterol (22OH; Avanti, 700058P) at the indicated concentrations. 22OH was serially diluted in medium containing 0.4% v/v anhydrous ethanol (vehicle; Commercial Alcohols, P016EAAN) to keep the concentration of vehicle constant. Cells were left to incubate for 48 h prior to addition of 10 µL per well of Cell Counting Kit 8 (CCK-8; ApexBio, K1018). Samples were seeded in triplicate and only the inner wells were used to reduce variation from evaporation. A colourmetric assay was used to monitor cell viability instead of a fluorescence-based assay to avoid any potential interference from the GFP and RFP expression from the iSLK.219 cells. After 1 h of incubation with CCK-8, absorbance was read at 450 nm with correction at 600 nm using a CLARIOstar Plus (BMG Labtech) plate reader. Values were imported into Prism (GraphPad) and CC50 values were calculated by fitting a non-linear curve to the data.

### Titring infectious KSHV

iSLK.219 cells were seeded into plates and the following day the medium was replaced in fresh medium containing 1 µg/mL doxycycline and serial dilutions of 22OH (prepared in medium containing 0.4% v/v anhydrous ethanol), type II water (dH_2_O), or 50 nM cerivastatin (prepared in dH_2_O; Sigma-Aldrich, SML0005). One day prior to harvest, 293A cells were seeded at 1.5 x 10^5^ cells/mL into Nunc MicroWell 96-Well Optical-Bottom Plates with Polymer Base (Thermo Fisher, 165305) that were previously coated with 5 µg/mL poly-L-lysine (Millipore-Sigma, P2658) in phosphate-buffered saline (PBS; Thermo Fisher, 10010023). Supernatants (5 µL) from the reactivated iSLK.219 cells were harvested 96 h post-reactivation, diluted with 95 µL medium, and then used to infect the 293A cells seeded by spinoculation (plates were centrifuged at 800 x *g* at 37°C for 90 min) then incubated for 24 h at 37°C. The cells were washed with PBS and incubated with 4% paraformaldehyde (PFA; Electron Microscopy Services, 15710) in PBS for 20 min. Cells were counterstained with 1:3000 Hoechst 33342 (Invitrogen, H21492) in PBS for 15 min at room temperature and GFP-positive (infected) cells were imaged using an Axio Observer widefield fluorescence microscope with a 10X objective and a Colibri 5 LED light source using automated image collection with Zen Blue v.3 (ZEISS). For Fig. 1C, KSHV-positive (GFP) cells and Hoechst stained muclei were quantitated using Imaris Microscopy analysis software (Oxford Instruments). For Fig. 1H, the raw images were converted to Tiff files and background corrected (rolling ball radius of 25 pixels) with Fiji (124). The modified images were segmented in Cellpose 3 (cyto3 pre-trained model (125) to identify regions of interest (nuclei or GFP). Titres were determined using the following formula: 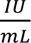 = ((− ln(1 − *Fraction infected*)) × 40000) ÷ 0.005 *mL* where 40000 is the approximate number of cells per 96-well and 0.005 mL is the input volume of virus containing supernatant. Graphing and statistically analysis (1-way ANOVA, Tukey test) were performed in Prism (GraphPad).

### Western blotting

iSLK.219 cells were reactivated with 1 µg/mL doxycycline and simultaneously treated with anhydrous ethanol (vehicle; Commercial Alcohols, P016EAAN) or 9 μM 22OH. Cells were harvested at 24, 48, and 72 h post-reactivation (iSLK.219) or hours post-infection (hpi; HSV). Cell lysates were harvested in 125 mM Tris-HCl (pH 6.8), 2% w/v sodium dodecyl sulfate (SDS), 10% v/v glycerol (2x Laemmli buffer) at the indicated times post-reactivation. Samples were homogenized using a 21-gauge needle or passed through a QIAshredder (QIAGEN, 79656) at 8000 x *g* for 2 min. Protein concentrations were determined using a DC Protein Assay Kit II (BIORAD, 5000112) prior to the addition of 100 mM dithiothreitol (DTT) and 0.005% w/v bromophenol blue and boiled at 95 °C prior to use. Samples with equal µg of protein were subjected to SDS-PAGE and immunoblotted for viral and host proteins using the indicated antibodies (antibody information can be found in Table 1).

**Table 1:** Antibodies for immunoblotting, immunofluorescence, and flow cytometry.

| *Primary antibodies* |  |  |  |  |
| --- | --- | --- | --- | --- |
| *Antigen* | *Supplier* | *Application* | *Catalog #* | *[Final]* |
| β-actin, HRP-conjugated (rabbit) | Cell Sig. | WB | 5125 | 1:1000 |
| ABCA1 (rabbit) | Novus | WB | NB400-105 | 1:1000 |
| HSV gG (mouse) | Virusys | WB | P1104 | 1:2500 |
| HSV ICP27 (mouse) | Abcam | WB | ab53480 | 1:1000 |
| HSV pUL31 (rat, (132)) | B. Banfield | IF | N/A | 1:500 |
| HSV pUL34 (chicken, (132)) | B. Banfield | IF | N/A | 1:500 |
| HSV pUS3 (rat, (133)) | B. Banfield | WB | ‘B’ | 1:1000 |
| KSHV ORF45 (mouse) | Thermo Fisher | WB | MA5-14769 | 1:1000 |
| KSHV ORF57 (mouse) | Santa-Cruz | FC | sc-135746 | 1:400 |
| KSHV ORF57 (mouse) | Santa-Cruz | IF | sc-135746 | 1:1000 |
| Lamin A/C (mouse) | Santa Cruz | IF | sc-7292 | ? |
| Lamin A/C (mouse) | Santa Cruz | WB | sc-7292 | 1:1000 |
| LXRα (mouse) | Abcam | WB | ab41902 | 1:1000 |
| *Secondary antibodies* |  |  |  |  |
| *Antigen* | *Supplier* | *Application* | *Catalog #* | *[Final]* |
| Anti-chicken IgG (H+L), Alexa Fluor ??? | Invitrogen | IF | A??? | ? |
| Anti-mouse IgG (H+L), Alexa Fluor 647 | Invitrogen | IF | A21463 | 1:1000 |
| Anti-rat IgG (H+L), Alexa Fluor ??? | Invitrogen | IF | A??? | ? |
| Anti-mouse IgG F(ab’)<sub>2</sub>, Alexa Fluor 647 | Invitrogen | FC | A21237 | 1:1000 |
| Anti-mouse IgG, HRP-linked | Cell Sig. | WB | 7076 | 1:5000 |
| Anti-rabbit IgG, HRP-linked | Cell Sig. | WB | 7074 | 1:5000 |
| Anti-rat IgG, HRP-linked | ThermoFisher | WB | 31470 | 1:5000 |

Abbreviations: Cell Sig., Cell Signaling; FC, flow cytometry; HRP, horseradish peroxidase; IF,
immunofluorescence; WB, western blotting.

### Cholesterol quantification

iSLK.219 cells were harvested in cold PBS at 48 h post-reactivation with anhydrous ethanol (vehicle) or 9 μM 22OH. Samples were centrifuged at 300 x *g*, flash frozen in liquid nitrogen, and stored at −80°C. Cholesterol was measured using an Amplex Red Cholesterol Assay Kit (Invitrogen, A12216) as per manufacturer’s instructions with the following changes: samples were re-suspended in 200 µL 0.1% w/v SDS in PBS and the cholesterol standard curve was diluted in 0.1% w/v SDS in PBS. A Student’s unpaired *t*-test was performed to determine statistical significance.

### DNase-protected extracellular KSHV genome quantification

iSLK.219 cells were reactivated with 1 µg/mL doxycycline and simultaneously treated with anhydrous ethanol (vehicle), 9 μM 22OH, or 500 µM phosphonoacetic acid (PAA, prepared in water; Sigma-Aldrich, 284270-10G). Supernatants were harvested at the indicated times (24, 48, 72, and/or 96 h) post-reactivation. Extracellular viral DNA was extracted using the DNeasy Blood and Tissue Kit (QIAGEN, 69506) with the following modifications: 500 μL of media was centrifuged at 5000 x *g* at 4 °C for 4 min, 180 μL of supernatant was mixed with 20 μL DNase I (3 mg/mL; Sigma-Aldrich, D4513) and incubated at 37 °C for 30 min, buffer mix was added to each sample (200 μL buffer AL (QIAGEN), 5 μg salmon sperm DNA solution (Invitrogen, 15632011), 100 pg luciferase vector (pGL4.26, Promega), and 20 μL proteinase K (Invitrogen, 4333793). Samples were incubated at 56 °C for 10 min. qPCR was performed using primers for *ORF26*, *β-actin*, and *luciferase*. All primers were purchased from Thermo Fisher and the sequences are provided in Table 2.

**Table 2:**
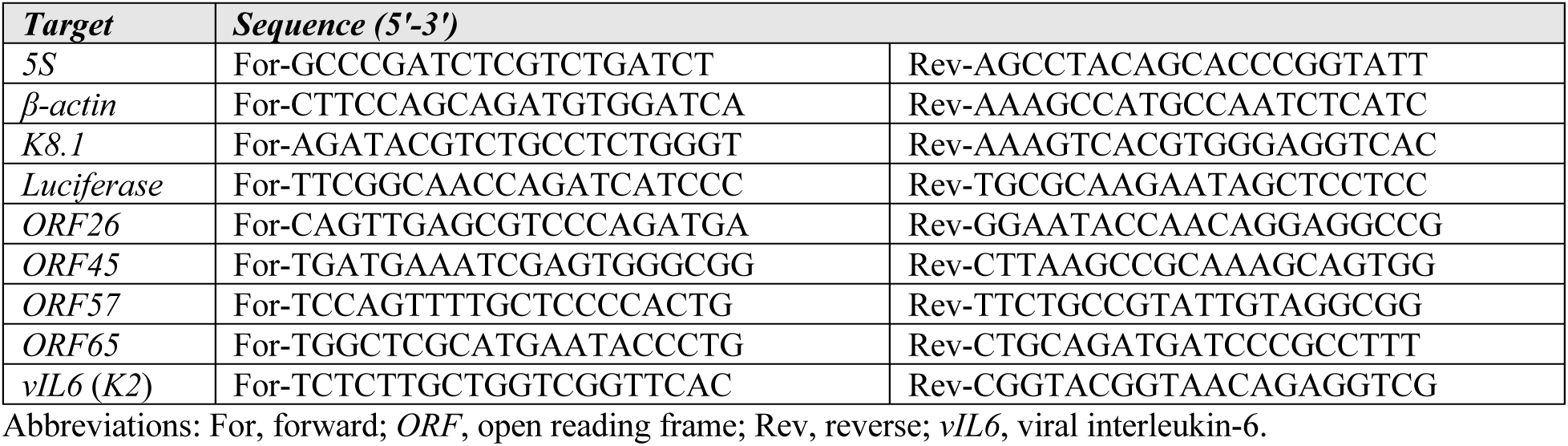
Primer sequences for qPCR.

### Fluorescence microscopy

For live cell microscopy, iSLK.219 cells were reactivated as previously described and treated with vehicle or 10 μM 22OH. GFP and RFP expression of iSLK.219 cells was imaged on an EVOS-FL Imaging System (Thermo Fisher) at 10X magnification. For immunofluorescence, cells were fixed in 4% PFA for 10 min at 37 °C and washed 3 times with PBS. Samples were permeabilized in 0.1% v/v Triton X-100 in PBS for 10 min at room temperature and washed with PBS before blocking in 1% v/v human serum (Sigma-Aldrich, 4522) in PBS for 1 h at room temperature. Samples were probed with mouse anti-KSHV ORF57 overnight at 4°C and anti-mouse IgG (H+L) Alexa Fluor 647 for 1 h at room temperature (see Table 1), both in 0.1% Triton X-100 in PBS. Nuclei were visualized by staining with DAPI (Invitrogen, D1306) for 5 min at room temperature. Samples were imaged on a LSM710 upright microscope (Zeiss) at 63X magnification with 1.4X zoom. Data were analyzed using ImageJ. A student’s *t*-test was performed to determine significance.

### Flow cytometry

iSLK.219 cells were seeded in 12-well plates at a density of 2.0 x 10^5^ cells/mL. The next day, the cells were treated with 10 μM 22-OH or anhydrous ethanol in the presence or absence of 1 μg/mL doxycycline. Compensation controls were generated by seeding 293T cells at 2.0 x 10^5^ cells/mL and the next day transfecting the cells using PEI MAX Transfection Grade Linear Polyethylenimine (Polysciences, 24765) and 1 ug of pEGFP-C1, pdsRed-Express, or pJLM1B-Puro-ORF57 plasmids. The iSLK.219 cells were harvested at 24 h (iSLK.219 and 293T cells), 48 h, or 72 h post-treatment using 5 mM ethylenediaminetetraacetic acid (EDTA) in PBS, followed by three washes with PBS, and fixation with 4% PFA in PBS. The cells were resuspended in 70% ethanol (prepared from Commercial Alcohols, P016EA95) and incubated at 4°C for 30 min. Then, the cells were washed with PBS and incubated with 1% human serum in PBS for 10 min at 4°C. Mouse anti-KSHV ORF57 in 1% human serum in PBS was added to the cells and incubated at 4°C for 20 min. The cells were then washed twice with PBS and incubated with secondary antibody solution containing goat anti-mouse IgG F(ab’)2 Alexa Fluor 647 secondary antibody in 1% human serum in PBS for 20 min at 4°C and then washed twice with PBS again. Finally, the cells were resuspended in 1% BSA/5 mM EDTA/PBS containing 0.1% sodium azide (Flow buffer) and fluorescence data was collected using a CytoFLEX (Beckman Coulter) equipped with 405 nm, 488 nm, and 638 nm lasers. Following acquisition, the data was analyzed using FCS Express v6.06.0042 (De Novo). Compensation was applied to control for EGFP/dsRed spillover and the gated single cells were evaluated for EGFP+ (KSHV+ cells), dsRed+ (reactivated KSHV+ cells), and ORF57+ (reactivated KSHV+ cells). Graphing and statistically analysis (2-way ANOVA, Holm-Šídák test) were performed in Prism (GraphPad).

### RT-qPCR

Viral and host RNA was extracted from iSLK.219 cells at 24, 48, and 72 hpi using the RNeasy Plus Mini Kit (QIAGEN, 74136). cDNA was synthesized using the Maxima H Minus First Strand cDNA Synthesis Kit (Thermo Fisher, K1652) using random hexamers. qPCRs were performed using 200 nM primers, 1:250 (final) diluted cDNA, and 1X GoTaq qPCR Master Mix (Promega, A6002). Amplifications were performed using a CFX Connect Real-Time PCR Detection System (Bio-Rad) and Bio-Rad CFX Manager 3.1 software. Data were analyzed using the ΔΔCt method. All primers were purchased from Thermo Fisher and the sequences are available in Table 2.

### Label-free mass spectroscopy

Latent iSLK.219 cells or cells reactivated with 1 µg/mL doxycycline were mock treated (anhydrous ethanol) or treated with 10 µM 22OH and were harvested at 48 h and 72 h post-reactivation then processed for label-free proteomics. For all experiments the cells were washed once with ice-cold PBS, scraped, pelleted at 500 x *g* for 5 min, then the washed once more in ice-cold PBS. Afterwards the cell pellet was snap-frozen in liquid nitrogen and stored at −80^°^C until processing. Protein was extracted from cell pellets using 150 μL 20mM Tris-Cl (pH 7.5), 150 mM KCl, 5 mM MgCl_2_, 0.5% (v/v) NP⁃40, 0.5% (w/v) sodium deoxycholate supplemented with 0.5X cOmplete EDTA-free protease inhibitor cocktail (Millipore Sigma, 4693132001). To support degradation of DNA/RNA, 10 μL of a nuclease solution (1 U/μL benzonase (Millipore Sigma, 70664)) was added to the solution and incubated at 24°C for 10 min. After nuclease digestion, 50 μL of a solution of 600 mM Tris-Cl (pH 7.5), 8% (w/v) SDS, and 10 mM DTT was added to the lysate and incubated at 95°C for 5 min. After cooling to 24°C, lysates were alkylated using 40 mM chloroacetamide (final concentration) and subsequently quenched with DTT. Protein concentration of the resulting lysate was measured using a Pierce BCA Protein Assay Kit (Thermo Fisher, 23227). Protein cleanup prior to digestion was carried out using an adapted version of the previously described SP3 protocol (126). Specifically, 1 μL of a mixture of Sera-Mag carboxylate-modified SpeedBeads (Millipore Sigma, GE45152105050250 and GE65152105050250, prepared as 100 mg/mL stock solution in water) was added to 50 μg of protein for each sample prior to four volumes of acetone. Mixtures were incubated on a Thermomixer at 37°C for 5 min at 800 RPM prior to centrifugation at 5000 x *g* for 5 min. After spinning, the supernatant was discarded and beads rinsed using 800 μL of 80% (v/v) ethanol solution with pipetting to resuspend the beads. Mixtures were centrifuged at 5000 x *g* for 5 min and the supernatant discarded prior to the addition of 1 μg of Trypsin/Lys-C Mix (Promega, V5072) in 100 μL of digestion solution (100 mM NH_4_HCO_3_, 1 mM CaCl_2_) and incubation at 37°C for 16 h in a Thermomixer with mixing at 800 RPM. After digestion, peptides were centrifuged at 12000 x *g* for 2 min and the supernatant recovered to a fresh tube containing 5 μL 10% (v/v) trifluoroacetic acid. To desalt peptides prior to mass spectrometry (MS) analysis, an HPLC system (Agilent 1290 Infinity II with a DAD module) equipped with a reversed-phase column (CORTECS T3 1.6μm, 2.1×50mm, Waters) was utilized. Specifically, peptides were injected into a column equilibrated at 4% mobile phase B (0.1% formic acid in acetonitrile), rinsed for 1.5 min at a flow rate of 1 mL/min, and subsequently eluted for 0.8 min at 80% mobile phase A (0.1% formic acid in water). Desalted peptides were dried in a SpeedVac centrifuge and reconstituted in 1% (v/v) formic acid in water at a concentration of 1 μg/μL based on the 214nm UV signal from the HPLC desalting step. Peptide samples were analyzed using a data-independent acquisition (DIA) acquisition routine on an Orbitrap Fusion Lumos mass spectrometer (MS) (Thermo Fisher). Samples were introduced to the MS using an Dionex UltiMate liquid chromatography (LC) instrument (Thermo Fisher) equipped with a trapping-analytical column setup. For injection, peptides were initially trapped using 95% mobile phase A on a 100 μm inner diameter x 3 cm length column packed in-house with 1.9 μm ReproSil-Pur C18 beads (Dr. Maisch, r119.aq.0001). Gradient elution of peptides was performed using a ramp of mobile phase B (80% v/v acetonitrile in 0.1% formic acid) on a 100 μm inner diameter x 25 cm length analytical column packed with 1.9 μm ReproSil-Pur C18 beads. For each injection, 1 μg of peptides were separated with an 80 min LC linear gradient from 5% to 11% mobile phase B in 0.5 min, and to 34% mobile phase B in 72.5 min at 400 nL/min, followed by rinsing and re-equilibration over the remaining 7 min. The analytical column outlet was coupled to a 20 μm inner diameter LOTUS electrospray tip (Fossil Ion Technology). The data presented was derived from three individual injections (80-minute acquisition time for each), with each acquisition covering analysis of a separate mass range (injection 1 = 430 - 550, 2 = 550 - 690, 3 = 690 – 930 m/z). The Orbitrap Lumos MS was globally set to use a positive ion spray voltage of 2200 V, an ion transfer tube temperature of 275°C, a default charge state of 3, and an RF Lens setting of 45%. The complete duty cycle of the acquisition method consisted of two MS1 scans and two sets of windowed MS2 scans (order MS1 - MS2 - MS1 - MS2). For the first, low mass range injection, the initial MS1 scan covered a mass range of 425-555 m/z at a resolution of 60000 with an AGC target of 4e^5^ (100%) and a max injection time set to ‘Auto’. The following set of DIA MS2 scans covered a precursor range of 430-550 m/z with an isolation window size of 4 m/z (0 m/z overlap) for a total of 30 scan events. Each scan used an HCD energy of 30% and covered a defined mass range of 200-1800 m/z at a resolution of 30000 with a normalized AGC target of 1000%, and the maximum injection time set to ‘Auto’. Loop control was set to ‘N’ with a value of 30 spectra. The next MS1 scan and following DIA MS2 used the same settings as the previous with the exception that the MS2 precursor mass range was set to 428-552 m/z to give a 2 m/z stagger with the previous scan windows. For the second, medium mass range injection, the scan settings were the same as for the low mass injection with the following changes: 1. MS1 scan range of 545-695 m/z; 2. DIA-MS2 precursor scan ranges of 550-690 m/z and 548-692 m/z; 3. Loop control number of spectra = 35. For the third, high mass range injection, the scan settings were the same as the others with the following changes: 1. MS1 scan range of 685-935 m/z; 2. DIA-MS2 precursor scan ranges of 690-930 m/z and 686-934 m/z; 3. DIA-MS2 isolation window of 8 m/z; 4. Loop control number of spectra = 30. All scan data were acquired in centroid mode. DIA-MS raw data were de-multiplexed using msconvert (ProteoWizard, version 3.0.23304-e66e43d, peakPicking = ‘vendor’, demultiplex = overlap only, 10ppm, SIM as spectra) and processed using DIA-NN (version 1.8.1) (127). Specifically, a representative human proteome fasta database (version 07/2024, 20,481 entries, includes non-human contaminants) concatenated with a KSHV fasta database manually generated from KSHV2.0 (75) was provided to DIA-NN along with all of the raw data in order to perform an initial spectral library generation step (default settings for library generation: min-pr-mz, 430; max-pr-mz, 930; min-pr-charge, 2; max-pr-charge, 4; MBR, disabled). Afterwards, a subset of raw files for each mass range were searched together against this generated spectral library using DIA-NN with ‘unrelated runs’ activated to derive appropriate mass error and scan window settings. The entire set of raw files for each mass range were then searched against the spectral library using the derived mass error and window settings with MBR enabled. Resulting report files were passed to the iq package (Lib.Q.Value = 0.01, Lib.PG.Q.Value = 0.01, Q.Value = 0.01, PG.Q.Value = 0.05) in R to generate estimates of protein abundance (128). Differential protein expression was determined using the R package DEqMS (129). Volcano plots were generated in Prism (GraphPad) and KRT73 (log_2_ FC −7.19, 48 h), STMN2 (log_2_ FC 6.86, 48 h), SIRPA (log_2_ FC −6.17, 72 h), COL18A1 (log_2_ FC −5.22, 72 h), and CTNNA2 (log_2_ FC 7.68, 72 h) are not shown in Fig. 2F. Gene Set Enrichment Analysis (GSEA) was performed with GSEA v.4.4.0 (86, 87) using the GSEAPreranked setting. Area-proportional Venn diagrams were generated using the plotter and editor available at bioinforx.com/apps/venn.php and edited in Affinity Designer.

### Transmission electron microscopy

iSLK-BAC16 cells were seeded in 10 cm dishes at ∼1.5 x 10^6^ cells per dish and reactivated and treated as described in the figure legend. Cells were harvested using 0.05% trypsin/EDTA (Gibco, 2533-054) treatment and centrifuged at 250 x *g* for 5 minutes. Samples were fixed for a minimum of 2 hours in 2.5% glutaraldehyde (Cedarlane, 16210(EM)) diluted in 0.1 M sodium cacodylate buffer (Cedarlane, 12300(EM)). Following fixation, samples were rinsed three times for at least 10 minutes each with 0.1 M sodium cacodylate buffer. Secondary fixation was performed using 1% osmium tetroxide (Ventron Alfa Divison) for 2 hours, followed by a brief rinse with distilled water. Samples were then placed in 0.25% uranyl acetate (Fisher Scientific) at 4°C overnight. Dehydration was carried out using a graded acetone series: 50% acetone for 10 min, 70% acetone 2 x 10 min, 95% acetone 2 x10 min, and 100% acetone 2 x 10 min, followed by 10 min in dried 100% acetone. Samples were then infiltrated with Epon-Araldite resin (Cedarlane, 13940(EM)) using a stepwise approach: 3:1 (dried 100% acetone to resin) for 3 h, 1:3 (dried 100% acetone to resin) overnight, and finally 100% resin for two 3 h incubations. Samples were embedded in 100% Epon-Araldite resin and cured at 60°C for 48 h. Ultrathin sections (∼100 nm thick) were obtained using a Reichert–Jung Ultracut E ultramicrotome equipped with a diamond knife and were placed on 300-mesh copper grids (cat#). Sections were stained with 2% aqueous uranyl acetate for 10 min, followed by two 5 min rinses with distilled water. Lead citrate staining was performed for 4 min followed by a quick rinse with distilled water. All grids were air-dried. Prepared samples were imaged using a JEOL JEM 1230 transmission electron microscope operating at 80 kV. Images were acquired using a Hamamatsu ORCA-HR digital camera. TEM experiments were performed in three biological replicates. Capsids were counted manually and assigned A-, B-, or C-capsid designations. In the event capsid type could not be assigned, a capsid was excluded from analysis.

### HSV plaque reduction assays

HSV-1 strain 17syn+ or HSV-2 strain 186 were the wild-type strains used. For propagation, Vero cells were infected at a multiplicity of infection of 0.05 and cell-associated virus was harvested once complete cytopathic effect was observed. Virus was liberated from cells following three freeze/thaw cycles and sonication. Lysate was cleared by centrifugation at 1500 x *g* for 5 min and the viruses titred on HeLa cells. HeLa cell monolayers were inoculated in serum free media with ∼150 plaque forming units of virus per well for 1 h with shaking every 15 min. Inoculum was removed and overlay (1% human serum in DMEM) was added, containing 0.1 μM, 1 μM or 10 μM of 22OH or anhydrous ethanol (vehicle). Four days post-infection, cells were fixed and stained with 0.5% crystal violet (prepared in 1:1 v/v methanol:water) for 15 min at room temperature and then washing with water. Plaque assays were imaged using a ChemiDoc Touch (BioRad). Plaque size and number were quantified using CellProfiler v3.0 (130) with a custom quantitation pipeline.

### Data management, software, and statistical analyses

22(*R*)-hydroxycholesterol chemical structure generated using PubChem (NIH-NLM). Data are shown as the mean ± standard error of the mean (SEM). Statistical significance was determined using the appropriate post-test (indicated in the corresponding figure legends or methods). All statistics were performed using GraphPad Prism v.7-v.10 (Dotmatics). Affinity Designer software v.1-2 was used for figure preparation. The mass spectrometry proteomics data have been deposited to the ProteomeXchange Consortium via the PRIDE (131) partner repository with the dataset identifier PXD068855.

## ACKNOWLEDGEMENTS

We thank Drs. Don Ganem (UCSF; Chan-Zuckerberg Biohub), Jae Jung (USC), Shou-Jiang Gao (Pitt), and David Lukac (Rutgers) for providing reagents. We thank the managers of the following Dalhousie University Facility of Medicine CORES facilities for their support: Biological Mass Spectrometry, Cellular & Molecular Digital Imaging, Electron Microscopy, and Flow Cytometry. This work was supported by a Research Nova Scotia Scotia Scholar doctoral award (to A.N.W.), a Canadian Institutes for Health Research (CIHR) doctoral award (to E.C.M.T.) and CIHR Operating Grants MOP-84554 (to C.M.) and PJT-162162 (to B.W.B.).

## AUTHOR CONTRIBUTIONS

Conceptualization: E.S.P., C.-A.R., B.A.D., and C.M.; Methodology: E.S.P., C.-A.R., A.N.W., E.C.M.T., J.K., A.L.-A.M., K.B., and B.A.D.; Formal Analysis: E.S.P, C.-A.R., A.N.W., and B.A.D.; Investigation: E.S.P., C.-A.R., A.N.W., E.C.M.T., J.K., A.L.-A.M., K.B., and B.A.D.; Writing – Original Draft: E.S.P., C.-A.R., B.A.D., and C.M.; Writing –Review & Editing: E.S.P., C.-A.R., A.N.W., E.C.M.T., J.K., A.L.-A.M., K.B., B.A.D., B.W.B., and C.M.; Visualization: E.S.P, C.-A.R., A.N.W., and B.A.D.; Supervision: B.W.B. and C.M.; Project Administration: E.S.P., B.A.D., and C.M.; Funding Acquisition: B.W.B. and C.M.

